# Butyrate enhances Clostridioides difficile sporulation *in vitro*

**DOI:** 10.1101/2023.04.27.538596

**Authors:** Michelle A. Baldassare, Disha Bhattacharjee, Julian D. Coles, Sydney Nelson, C. Alexis McCollum, Anna M. Seekatz

**Author notes:** Corresponding Author. Life Sciences Building 157A, 190 Collings St, Clemson, South Carolina – 29634, United States of America. authors contributed equally (Michelle A. Baldassare, Disha Bhattacharjee).

## Abstract

Short chain fatty acids (SCFAs) are products of bacterial fermentation that help maintain important gut functions such as the intestinal barrier, signaling, and immune homeostasis. The main SCFAs acetate, propionate, and butyrate have demonstrated beneficial effects for the host, including importance in combatting infections caused by pathogens such as *Clostridioides difficile*. Despite the potential role of SCFAs in mitigating *C. difficile* infection, their direct effect on *C. difficile* remains unclear. Through a set of *in vitro* experiments, we investigated how SCFAs influence *C. difficile* growth, sporulation, and toxin production. Similar to previous studies, we observed that butyrate decreased growth of *C. difficile* strain 630 in a dose-dependent manner. The presence of butyrate also increased *C. difficile* sporulation, with minimal increases in toxin production. RNA-Seq analysis validated our experimental results, demonstrating increased expression of sporulation-related genes in conjunction with alternative metabolic and related *C. difficile* regulatory pathways, such as the carbon catabolite repressor, CcpA. Collectively, these data suggest that butyrate may signal alternative *C. difficile* metabolic pathways, thus modifying its growth and virulence to persist in the gut environment.

**IMPORTANCE:** Several studies suggest that butyrate may be important in alleviating gut infections, such as reducing inflammation caused by the healthcare-associated *Clostridioides difficile*. While studies in both animal models and human studies correlate high levels of butyrate with reduced *C. difficile* burden, the direct impact of butyrate on *C. difficile* remains unclear. Our study demonstrates that butyrate directly influences *C. difficile* by increasing its sporulation and modifying its metabolism, potentially using butyrate as a biomarker to shift survival strategies in a changing gut environment. These data point to additional therapeutic approaches to combat *C. difficile* in a butyrate-directed manner.

## INTRODUCTION

*Clostridioides (Clostridium) difficile* is an anaerobic, gram-positive bacterium of serious concern, causing nearly half a million infections and 30,000 deaths each year in the United States (1). *C. difficile* infection (CDI) causes inflammation and colitis in the gut, with symptoms ranging from diarrhea to pseudomembranous colitis and megacolon in extreme cases(2). Recurrence occurs in up to 30% of individuals resulting in higher patient mortality and increased healthcare costs making CDI an important public health threat (3). Risk factors for CDI include advanced age, pre-existing gastrointestinal issues, immunocompromised status, and antibiotic exposure, making CDI highly prevalent in healthcare-associated environments (4).

The gut microbiota, the microbial community residing in the gastrointestinal tract, provides colonization resistance against *C. difficile* within the colon. In most healthy individuals, contact with metabolically inert *C. difficile* spores from the environment does not result in disease. However, antibiotics and other environmental perturbations have been demonstrated to disrupt the microbiota (5, 6), allowing for *C. difficile* spores to colonize the gut environment and produce toxins, after germinating into its vegetative state (7). During initial colonization, removal of microbes that transform cholic acid and conjugated derivatives (primary bile acids), known to induce *C. difficile* germination (8), into deoxycholic acid (secondary bile acids), known to reduce *C. difficile* growth (9, 10), have been correlated with the development of CDI (5). Once colonized, *C. difficile* can use a variety of metabolic approaches to persist in the gut. For instance, the ability of *C. difficile* to use a multitude of carbohydrate sources for growth (11) as well as the ability to use amino acids for growth via Stickland fermentation (12, 13), likely supporting its ability to colonize multiple nutritional niches once competing organisms have been removed from the gut.

Metabolic flexibility of *C. difficile* is also connected to virulence mechanisms. The nutritional regulators CodY (14, 15) and CcpA (11) are known to decrease toxin production and sporulation by sensing nutrient deprivation or carbohydrate availability, respectively. Other regulators include PrdR, a proline regulator shown to be required for *C. difficile* growth (16), and Rex, a global redox sensing regulator, both of which influence toxin and spore production (17, 18). The main toxins produced by *C. difficile* include TcdA and TcdB, expressed by the *tcdA* and *tcdB* genes located on the PaLoc, or pathogenicity locus (19). Production of TcdA/B is regulated by the additional PaLoc genes *tcdR* and *tcdC*, which are further regulated by global transcriptional networks that respond to environmental cues (20). Sporulation, also a consequence of colonization, is controlled by master transcriptional regulator, *spo0A,* and the subsequent intricate regulation, leading to persistence of *C. difficile* in a compromised environment further ravaged by therapeutic antibiotics.

The high recurrence rate observed for CDI following standard antibiotic treatment has led to interest in developing microbial-mediated treatments that aim to recover colonization resistance against *C. difficile* (21). In addition to bile acid transformation, other microbiota-mediated mechanisms hypothesized to control *C. difficile* infection include short chain fatty acids (SCFAs), which are fermentation end-products produced by select microbes that are generally regarded as beneficial (22, 23). In particular, the SCFA, butyrate, has been correlated with recovery from CDI following treatment with fecal microbiota transplantation (FMT) (22, 23), which aims to restore microbial functions that provide colonization resistance against *C. difficile.* Butyrate has also been demonstrated to decrease *C. difficile* growth *in vitro* (24), as well as alleviate toxin-based inflammation in an animal model of CDI without directly reducing *C. difficile* burden (25). Yet, the mechanism by which butyrate might control *C. difficile* pathogenesis is relatively undefined.

This study sought to identify how SCFAs might directly influence *C. difficile* pathogenesis. Using an *in vitro* platform, we observed that in addition to attenuating growth, butyrate and propionate, increased sporulation of *C. difficile* strain 630. Butyrate’s effects were dependent on the nutritional environment, suggesting its effects might be metabolically regulated. RNA-Seq validated the observed experimental effects of butyrate, and further identified involvement of the major regulators CcpA and Spo0A, as well as a putative carbon-starvation gene, CstA, in butyrate-dependent control of C. *difficile*. Collectively, these results point to additional considerations in targeting butyrate as a therapeutic strategy to prevent or treat *C. difficile*.

## MATERIALS AND METHODS

### In vitro growth of *C. difficile*

In an anaerobic chamber (Coy), a 10^2^ dilution of the noted spore stock (*C. difficile* strain 630 (ATCC BAA-1382), VPI10463 (ATCC 43255-FZ), and R20291 (26)) prepared as described previously (27) was plated on pre-reduced taurocholate-cefoxitin-cycloserine-fructose agar (TCCFA) plate (28) and incubated overnight at 37°C. A single colony was inoculated into 5 mL of 1X of brain heart infusion broth supplemented with 5 g/liter yeast extract and 0.1% l-cysteine (BHI) (29), in biological triplicates per treatment group, and incubated at 37°C. After 18 hours of growth, tubes were centrifuged at 1500 x g for 10 minutes. After discarding the supernatant, each pellet was resuspended in 1 mL of 2X (twice concentrated) BHI. For each technical triplicate (three per biological replicate), 250 μL of the resuspended pellet was added into 5 mL of 2X BHI. In a 96-well plate, 100 μL of the prepared inoculum was added into each of the wells, making the final volume 200 μL. For negative control wells (no *C. difficile*), 2X BHI was added instead.

To test the effect of acetate, propionate, and butyrate on the growth of the prepared inoculum above, previously prepared 500 mM SCFA stocks of each (frozen until use) were diluted to a final working concentration of 50 mM in the anaerobic chamber a day prior. A 96-well plate was set up to test 5 mM and 25 mM acetate, propionate, and butyrate from 50 mM concentration. The plate was then placed into a Sunrise plate reader (Tecan) for 24 hours at 37°C, where optical density (OD_600_) was measured every 15 minutes. After 24 hours, colony-forming units (CFUs) were assessed, as described below. The plate was covered in parafilm and placed in the −80°C freezer for storage for toxin assays.

For assessment of *C. difficile* growth at multiple timepoints, two replicate 96 well plates were prepared simultaneously, one placed in the plate reader and one in the incubator at 37°C to acquire matched OD_600_ and CFU measurements. At 6, 12, 18, and 24 hours, three wells per treatment were sampled and diluted for calculating colony forming units (CFUs). CFUs were quantified for each corresponding treatment or timepoint. For three biological triplicates per treatment, bacterial growth was serially diluted from 10^-1^ to 10^-5^. CFUs/mL was determined by the log_10_ of colonization, determined by number of colonies x dilution of plate counted x dilution factor. For assessment of spores, 20 μL of the sample was added to PCR tubes and the tubes were heated for 20 minutes at 65°C to kill off vegetative cells. After heating, the samples were plated on TCCFA plates using the same dilutions as above.

For assessing butyrate dose response on *C. difficile* growth, a working stock of butyrate from a previously frozen 500 mM stock was prepared initially. A 96-well plate was set up as described above, comprising of *C. difficile* inoculum in BHI supplemented with final concentrations of 5 mM, 10 mM, 25 mM, and 50 mM butyrate in the wells.

To investigate the effect of pH, BHI was prepared with a final pH of 6.2, 7.2, and 8.0. Using a pH probe (Mettler Toledo FiveEasy Plus), pH values were confirmed before autoclaving, after autoclaving, and after preparing growth conditions (including SCFA addition).

For assessing the effect of butyrate on *C. difficile* growth under single carbohydrate sources, 96-well plates were prepared as described above except using 2X *C. difficile* Minimal Media (CDMM) (30, 31) supplemented with 1% of the indicated sugar instead of BHI. A stock concentration of 2% weight by volume of the carbohydrates (glucose (32), fructose (33), lactose (34), maltose (35), trehalose (36), cellobiose (37), sucrose (38), mannitol (39), mannose (32), and raffinose (5)) was prepared and final concentration of 100 μL was added into the experimental plate. Growth of *C. difficile* was assessed using OD_600_ as described above.

### *C. difficile* toxin assay

This protocol was adapted from Theriot et al (40). Briefly, filtered media consisting of Dulbecco’s Modified Eagle Medium (DMEM) (Gibco DMEM #11965-092), with 5% fetal bovine serum (FBS) (Fisher Gibco™ Fetal Bovine Serum, Qualified, Heat Inactivated, US Origin #16-140-071), and PenStrep (Life Technologies Gibco™ Penicillin Streptomycin 5000 U/ml (Penicillin 5000 U/mL; Streptomycin 5000 μg/mL) #15070063) was used to propagate Green African monkey kidney epithelial (Vero) cells to confluence within a 96-well plate, at a density of 10^3^ cells / well based on the number of viable cells observed in a 1:1 mixture of cell suspension in trypan blue. The seeded plate was incubated for an hour at room temperature before placed at 37°C overnight. Prior to addition of samples for toxin assessment, old media was replaced with fresh media. For the toxin assay, cell growth samples (i.e., spent media from *C. difficile* growth assays in BHI supplemented with the indicated SCFAs) were filtered with a 20 μm filter. A dilution plate was prepared serial dilutions of cell culture filtrate, diluted up to 10^-6^. For a positive control, 0.01 μg/μL Toxin A (Invitrogen #10977-015) was added to PBS. The seeded plate was incubated for 40 minutes at room temperature to allow for anti-toxin activity, then placed at 37°C overnight. Toxin activity was determined by the presence of confluence (>75% confluent) under a microscope for the last dilution of each sample. The amount of toxin was quantified as log_10_(antitoxin dilution factor x Vero cell dilution factor x last dilution with cell rounding x initial PBS dilution).

### Assessment of spore production using phase contrast

Spore stock was streaked anaerobically onto TCCFA and incubated at 37°C for 24 hours. An isolated colony was inoculated into BHI for 18 hours at 37°C (in triplicates). Tubes were centrifuged at 1500 x g for 10 minutes. After discarding the supernatant, the pellet was resuspended with 1 mL of 2X 70:30 sporulation media (41). For each replicate, resuspended pellet was added into 2X 70:30 media at 1:200. For conditions supplemented with 25mM butyrate, a working stock of 50 mM was prepared as described above. A 96-well plate was set up to test sporulation efficiency of 70:30 media with or without butyrate, with each well for a final volume of 200 μL. The 96 well plate was then placed into the Tecan plate reader for 24 hours at 37°C to assess growth as described above.

After 24 hours, the *C. difficile* grown in 70:30 and butyrate were added together in separate tubes and centrifuged for 30 seconds at 13,000 rpm at room temperature. The cells were then resuspended in 25 μL of BHI. A microscope slide was then prepared for both conditions by adding 5 μL of the resuspended culture to a microscope slide and adding a coverslip. Phase contrast images were captured at 100X on Ph3 on a phase contrast microscope (Leica DM750 fitted with Leica ICC50W camera). Sporulation efficiency was calculated through the following equation: (spores)/(vegetative cells + spores) x 100 (41). At least 1000 cells were counted to get an accurate efficiency. Original cell density was also calculated using a spectrophotometer.

### RNA extraction

*C. difficile* was grown with or without butyrate (25 mM) as described above. *C. difficile* culture was collected and immediately frozen for RNA extraction at points representing early-log (∼ 0.2 OD_600_) and late-log (∼ 0.5 OD_600_. For growth in BHI without butyrate, culture was collected at seven (early-log) and ten hours (late-log); for growth in BHI supplemented with butyrate, culture media was collected at ten (early-log) and thirteen (late-log) hours. Before RNA extractions, samples were thawed and centrifuged at 10,000 rpm for 10 minutes at 4°C. The supernatant was then discarded, and the pellet was resuspended in 1 mL of 1:100 BME/water dilution. The samples were then centrifuged at 14,000 rpm for 1 minute at 4°C. The supernatant was discarded, and the cell pellet was resuspended in 1 mL of Trizol. The samples were incubated at room temperature for 15 minutes. The samples were then centrifuged at 5,000 rpm for 15 minutes at 4°C. All further extraction steps required the aqueous phase (42). The Zymo Direct-zol RNA Miniprep Plus extraction kit (Zymo Research #R2071) was used to extract the RNA. Qubit (ThermoFisher Scientific #Q33230) was then performed to confirm the concentration of RNA before sending off for sequencing.

### RNA-Seq and data analysis

RNA collected from growth experiments above was sent to the Microbial Genome Sequencing Center (MiGS, Pittsburgh (www.migscenter.com)) for Illumina sequencing (NextSeq 2000). Raw reads were quality checked and adapter-trimmed using Trim-galore (Supplementary Table S1) (43). Metaphlan was used to identify relative species abundance of sequence reads (44). Sequences were aligned using RSEM (45) to the *C. difficile* strain 630 reference GCA_000009205.2 under the accession number AM180355 (PRJNA78) (46, 47). FeatureCounts from subread was utilized to quantify reads (48). The DESeq2 package (49) was used then to analyze the differential expression, identifying genes that were significant. The RNA-Seq data in R using ggplot2 (50) was used to visualize results. Gene set enrichment and KEGG enrichment of the top ranked genes were analyzed using clusterprofiler with Wald’s statistic (51). All codes used to analyze can be accessed here:

### qRT-PCR

Following RNA extractions, cDNA of the samples was made following the NEB M-MuLV Reverse Transcriptase protocol (NEB #M0253). Qubit was used to assess the concentration of cDNA in the samples. Primers used included *rpoC* (housekeeping gene) (FP: CCAGTCTCTCCTGGATCAACTA, RP: CTAGCTGCTCCTATGTCTCACATC) (52), *tcdR* (FP: TTATTAAATCTGTTTCTCCCTCTTCA, RP: AGCAAGAAATAACTCAGTAGATGATT) (53), *tcdC* (FP: GAGCACAAAGGGTATTGCTCTA, RP: AAATGACCTCCTCATGGTCTTC) (54), *codY* (FP: CTCATCTTCTATAACTGAACTGTCTTGAAC, RP: TTTGATTTACTGGCCGGAGCATTG) (15), *ccpA* (FP: TCTTGTTCAACTATCCATGAAATCATAAC, RP:

AAATGGGATAGAAGAGGTTGCTAAA) (55), and *rex* (FP: TGGTGGATTTGGACAACAAGGA, RP: TGCTCCTACAAGAACTGCGT) (generated for this study). Reactions were made using iQ SYBR Green Supermix (BioRad #1708880) using manufacturer’s directions with a final cDNA concentration of 5 ng. Samples were run in triplicate with biological replicates using manufacturer’s directions. Expression levels were quantified/normalized using the housekeeping gene *rpoC* (42). The 2^-ΔΔCT^ method was used to calculate relative expression fold change between the control (*C. difficile* + BHI) and treatment groups (*C. difficile* + BHI+ butyrate 25 mM) in the genes of interest (*tcdR, tcdC, codY, ccpA, and rex*) compared to the housekeeping gene (*rpoC*) (56).

### Statistical Analysis

Significance was determined using one-way ANOVA for area under the curve (calculated using growthcurver in R) followed by post-hoc Dunnett’s test (using DescTools in R) for the growth curves. The significance on plate counts and toxin activity at different time points was tested using one-way ANOVA followed by Dunnett’s test. Significance on the spore efficiency was tested using Bonferroni pairwise t-test. Significance on the transcriptomics was calculated by DESeq2 using Wald’s test.

### Data Availability

All code used to analyze data are available at https://github.com/SeekatzLab/C.difficile-butyrate. The raw reads from RNASeq are available under BioProject number: PRJNA955248 (BioSamples SAMN34190971 – SAMN34190982).

## RESULTS

### Butyrate decreases *C. difficile* growth

To assess the effect of the predominant SCFAs on *C. difficile* growth *in vitro*, we grew different strains of *C. difficile* (630, VPI10463, R20291) in the presence of low (5 mM) and high (25 mM) concentrations of acetate, propionate, and butyrate supplemented in the rich medium, BHI (Figure 1 and Supplementary Figure S1). We observed significantly decreased growth of *C. difficile* strain 630 in the presence of both butyrate and propionate (5 and 25 mM) concentrations (Dunnett’s test on area under the curve (AUC), p < 0.001 and p < 0.05) using OD_600_ measurements (Figure 1A). At 24h, we observed significantly decreased growth (Dunnett’s test, p < 0.01) of *C. difficile* strain 630 using CFU enumeration in 25 mM butyrate-supplemented BHI (Figure 1B). We also observed decreased growth of *C. difficile* strain 630 using CFU enumeration at 6 and 24 hours in butyrate- and propionate-supplemented BHI (Dunnett’s test, p < 0.001) (Figure 1C). While a previous study have demonstrated butyrate-induced growth defects across multiple *C. difficile* strains (57), we did not observe significant differences in growth with any SCFA for two additional strains, *C. difficile* VPI 10463 and R20291 (Supplementary Figure S1). Decreased growth was also dose-dependent, as increased concentrations of butyrate up to 50 mM concentrations increasingly impaired *C. difficile* strain 630 growth, with higher butyrate concentrations starting at 10 mM butyrate (Figure 1D). To preclude the possibility that butyrate’s impact on *C. difficile* growth was pH-dependent, media with and without butyrate was adjusted to 7.2, as well as tested at pH 6.2 and 8. Significant growth decrease in the presence of 25 mM butyrate was still observed at pH 6.2 and 7.2 levels, with no significance at a more basic pH of 8.0 (Supplementary Figure S2). Given our strain-specific results, we focused on *C. difficile* strain 630 for the remaining experiments.

**Figure 1:**
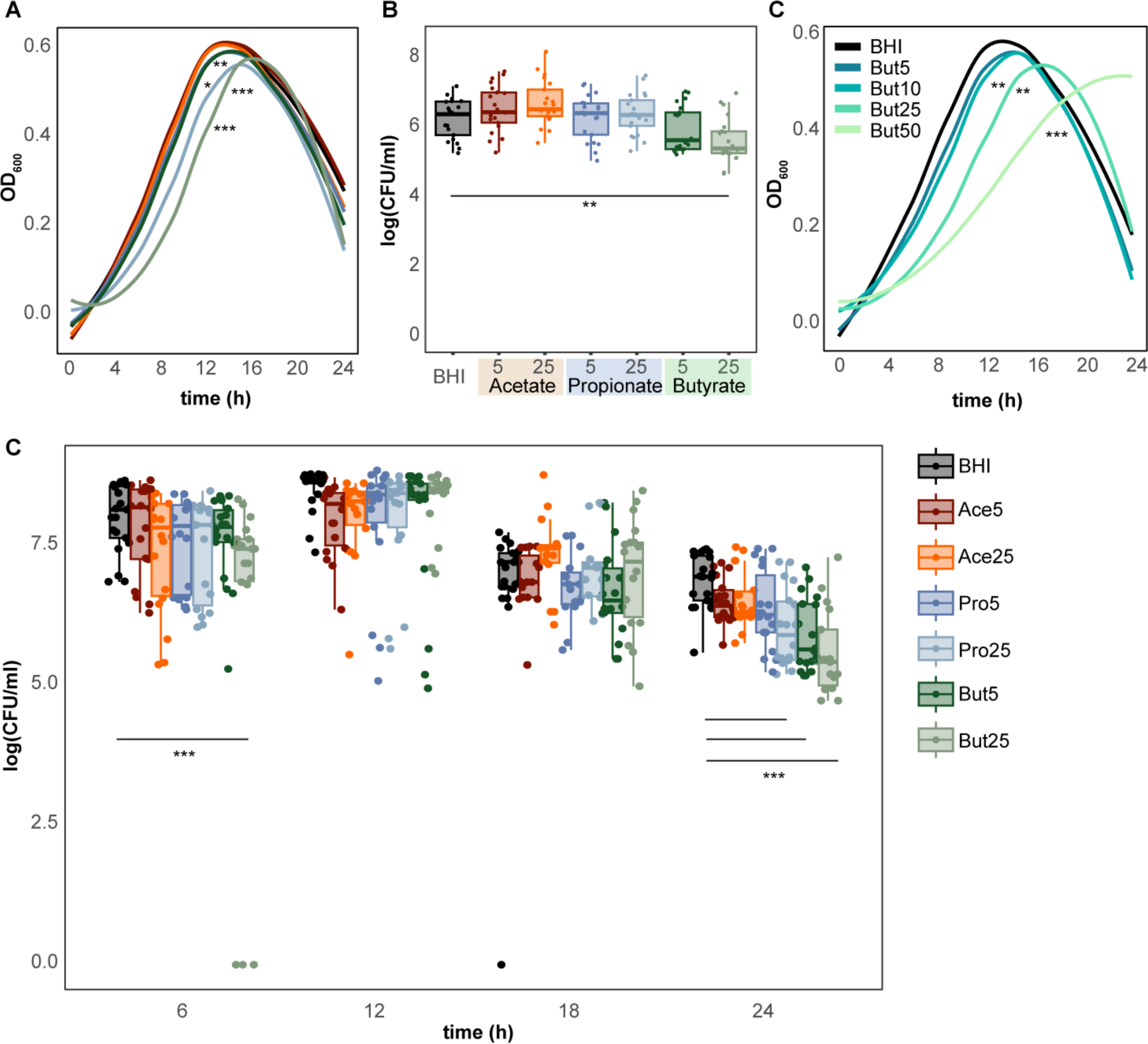
Butyrate inhibits the growth of *C. difficile* strain 630. **A)** Growth curves (OD_600_) over 24 hours. **B)** Colony-forming units (CFUs) after 24 hours and **C)** at 6, 12, 18 and 24 hours of growth in BHI supplemented with 5 and 25 mM acetate, propionate, or butyrate. **D)** Growth curves (OD_600_) over 24 h in BHI supplemented with increasing concentrations of butyrate (0, 5, 10, 25, 50 mM). Statistical significance calculated using Dunnett’s test: *p-value < 0.05; **p-value < 0.01; ***p-value < 0.001.

To test whether butyrate’s ability to modulate *C. difficile* growth is dependent on its metabolic environment, we grew *C. difficile* with or without butyrate in minimal media (CDMM) supplemented with different single carbohydrate sources known to support *C. difficile* growth (Figure 2) (58, 59). We observed decreased growth of *C. difficile* with butyrate in CDMM supplemented with 1% lactose and raffinose only (Dunnett’s test, p < 0.05) (Figure 2). The addition of butyrate did not significantly influence growth of *C. difficile* in CDMM supplemented with cellobiose, maltose, or trehalose. Surprisingly, butyrate increased the growth of *C. difficile* in CDMM supplemented with fructose, mannose and mannitol (Dunnett’s test, p < 0.01), and trended towards increase in growth in the presence of glucose and sucrose (not significant). These results suggest metabolism-dependent impacts of butyrate on *C. difficile* growth.

**Figure 2:**
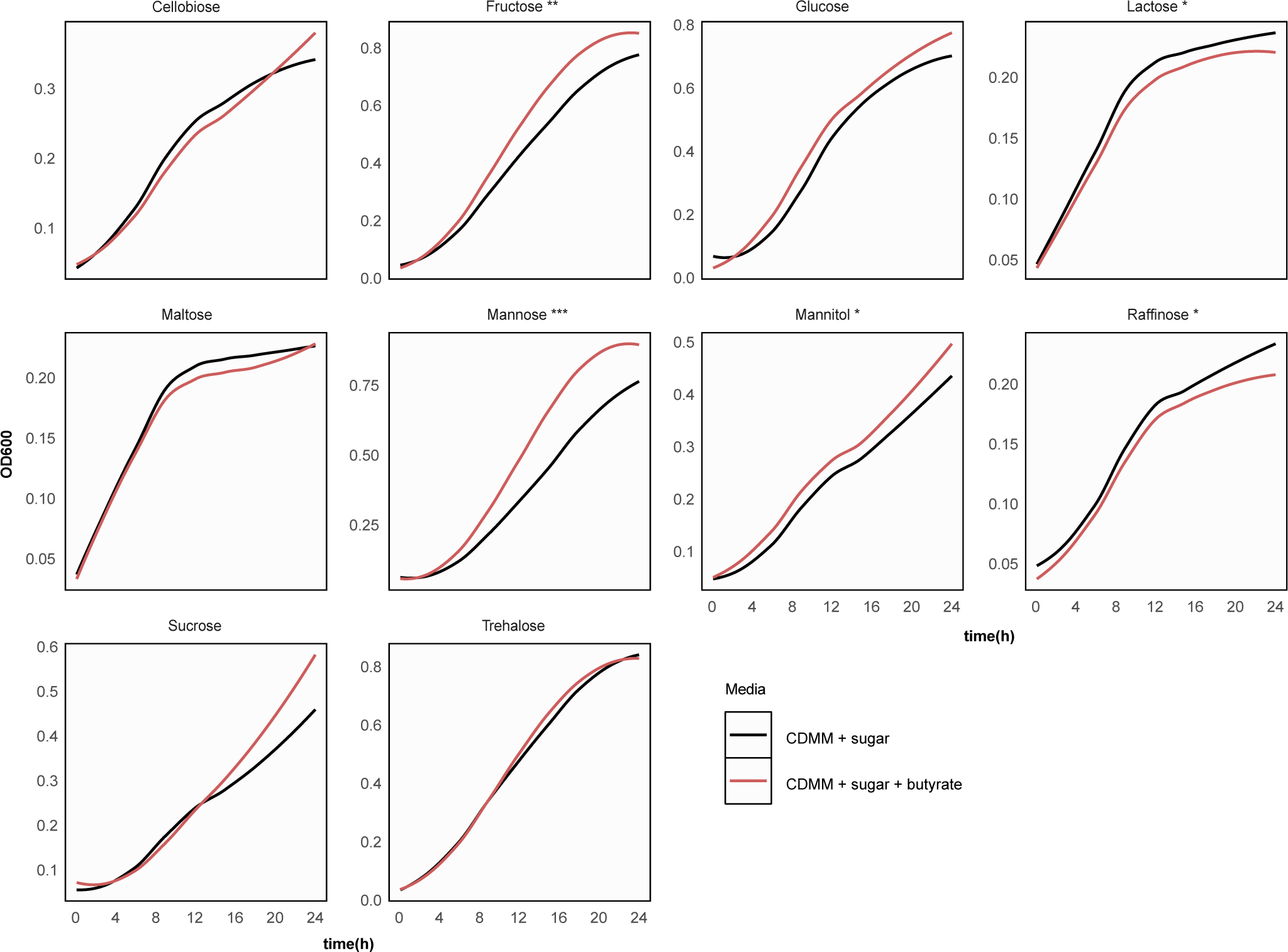
Butyrate-induced inhibition of *C. difficile* growth is dependent on the metabolic environment. Growth curves of *C. difficile* strain 630 (OD_600_) over 24 hours in minimal media (CDMM) in presence of a single sugar supplemented with (red) and without 25 mM butyrate (black). Statistical significance calculated using Dunnett’s test: *p-value < 0.05; **p-value < 0.01; ***p-value < 0.001.

### Butyrate increases toxin and spore production in *C. difficile*

We next assessed the impact of SCFAs on *C. difficile* toxin and spore production, which typically occur during stationary stage (60). We assessed toxin production in the presence of SCFAs using an *in vitro* cell rounding assay (61). After 24 hours of growth, we observed increased toxin by *C. difficile* strain 630 in the presence of all SCFAs, compared to BHI alone (Figure 3A, Dunnett’s Test, p < 0.001). All SCFA’s significantly increased toxin activity as early as 6 to 12 h in strain 630 (Supplementary figure S3C, Dunnett’s test, p < 0.05, 0.01, 0.001). Despite minimal effects on growth, SCFAs increased toxin activity in *C. difficile* strains VPI10463 and R20291 (Supplementary figure S3A and S3B, Dunnett’s test p < 0.05), which typically produce more toxin than *C. difficile* strain 630 (62). Enhanced and significant toxin activity was also observed at increasing butyrate concentrations starting at 12 h as low as 5 mM, and all concentrations except at 10 mM at 24 h (Figure 3B, Dunnett’s test, p < 0.01). To further validate these observations, we used qRT-PCR to assess the expression of *tcdR* and *tcdC*, the positive and negative regulators of *C. difficile* toxins located on the PaLoc (63), during early (∼ 0.2 OD_600_) and late-log (∼ 0.5 OD_600_) growth of *C. difficile* with or without butyrate (Figure 3C). In the presence of butyrate, *tcdC* expression was increased during both early- and late-log growth, whereas *tcdR* expression was decreased during both late- and early-log growth, albeit not significantly.

**Figure 3:**
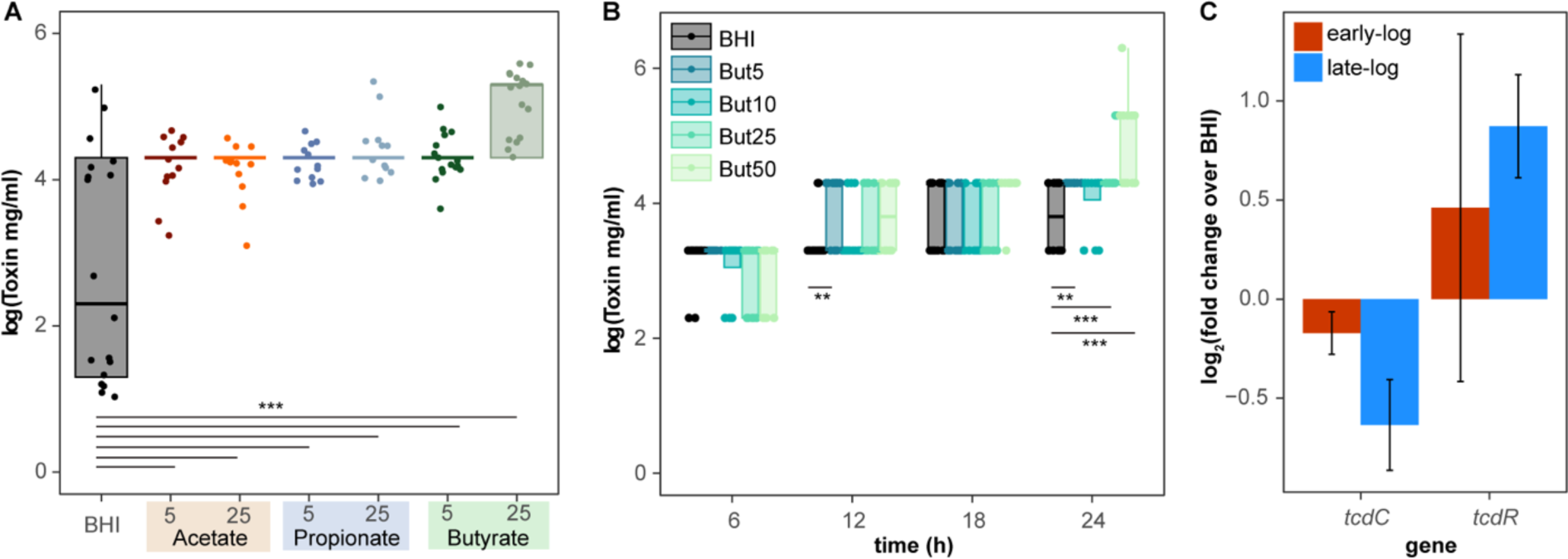
Short chain fatty acids increase *C. difficile* toxin production. Toxin activity of *C. difficile* strain 630, as measured by an *in vitro* cell assay **A)** after 24 hours of growth in BHI supplemented with 5 and 25 mM acetate, propionate, and butyrate, and **B)** at 6, 12, 18 and 24 hours in BHI supplemented with increasing concentrations of butyrate (0, 5, 10, 25, 50 mM). **C)** Log fold change of RNA expression in *tcdC* and *tcdR* (in culture with 25 mM butyrate over culture grown in just BHI) measured at early- and late-log growth using qRT-PCR. Statistical significance calculated using Dunnett’s test, *p-value < 0.05; **p-value < 0.01; ***p-value < 0.001.

To assess spore CFUs, *C. difficile* 630 cultures were heated at 65°C for 20 minutes to kill the vegetative cells prior to plating anaerobically on TCCFA. Compared to BHI alone, we observed significantly higher spore counts in the presence of 25 mM propionate and butyrate after 24 h (Dunnett’s test, p < 0.05, 0.01) (Figure 4A). Significantly increased spores were also observed as early as 6 h and later at 18 h specifically in presence of butyrate, even as low as 5 mM (Figure 4B Dunnett’s test, p < 0.05). Additionally, more spores were observed at increasing butyrate concentrations compared to BHI alone (Figure 4C, Dunnett’s Test, p <0.001). The ability of butyrate to increase spore production was also observed using phase contrast microscopy. Using a modified sporulation assay and phase contrast microscopy (41), we calculated the sporulation efficiency with and without butyrate after 24 hours of growth (Figure 4D, 4E, 4F). We observed higher sporulation efficiency in the presence of butyrate (13.26%) compared to BHI alone (3%). (Figure 4F, Welch’s two sample test, p < 0.001).

**Figure 4:**
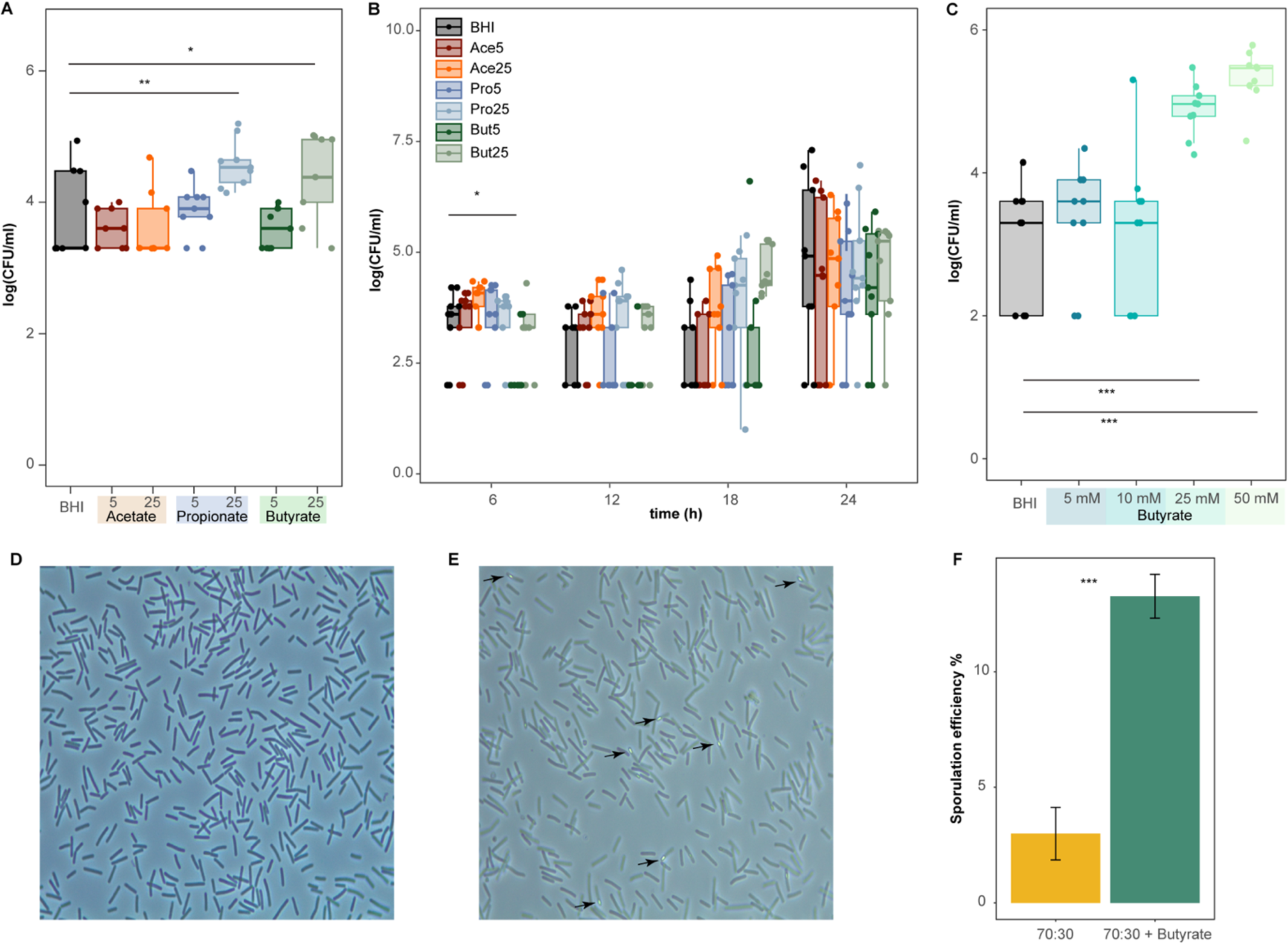
Butyrate increases *C. difficile* spore production. **A)** Colony-forming units (CFUs) of *C. difficile* strain 630 spores **A)** after 24 hours of growth and **B)** throughout growth at 6, 12, 18, 24 hours in BHI supplemented with 5 and 25 mM acetate, propionate, and butyrate, or **C)** after 24 hours of growth in BHI supplemented with increasing concentrations of butyrate (5, 10, 25, and 50 mM). Cultures were collected at indicated timepoints and heated at 65°C for 20 minutes to kill off vegetative cells, reflecting spore CFUs. Phase contrast (representative) of *C. difficile* strain 630 cells grown for 24 hours in **D)** 70:30 media alone or **E)** supplemented with 25 mM butyrate. **F)** Sporulation efficiency calculated over 1000 cells in 70:30 media with or without 25 mM butyrate. Welch’s two-sample test, *p-value < 0.05; **p-value < 0.01; ***p-value < 0.001.

### Butyrate induces the expression of genes related to spore production and alternate metabolic pathways

To determine global expression changes induced by butyrate, we conducted RNASeq of *C. difficile* strain 630 grown in BHI with and without 25 mM butyrate at both early-(∼ 0.2 OD_600_) and late-log (∼ 0.5 OD_600_) growth, and mapping the sequencing reads to *C. difficile* strain 630 (GCA_000009205.2; BioProject: PRJNA78). Nonmetric multidimensional scaling (NMDS) of all identified genes demonstrated clustering of samples based on the presence of butyrate (Figure 5A). Initial gene set enrichment analysis (GSEA) on genes expressed during early- or late-log growth identified differential pathways with or without butyrate (cutoff > log2 fold change, with adjusted p < 0.05 calculated with Wald’s test). During early-log growth, genes related to the phosphotransferase system and sucrose and starch metabolism were over-represented in the presence of butyrate, whereas genes related to nucleotide metabolism, aminoacyl-tRNA biosynthesis and ribosomal functions were under-represented when the genes are in decreasing order of log2 fold change (Figure 5B). During late-log growth, genes related to peptidoglycan biosynthesis were over-represented in the presence of butyrate, whereas genes related to secondary metabolites, amino acid, and carbon metabolism were under-represented (Figure 5C).

**Figure 5:**
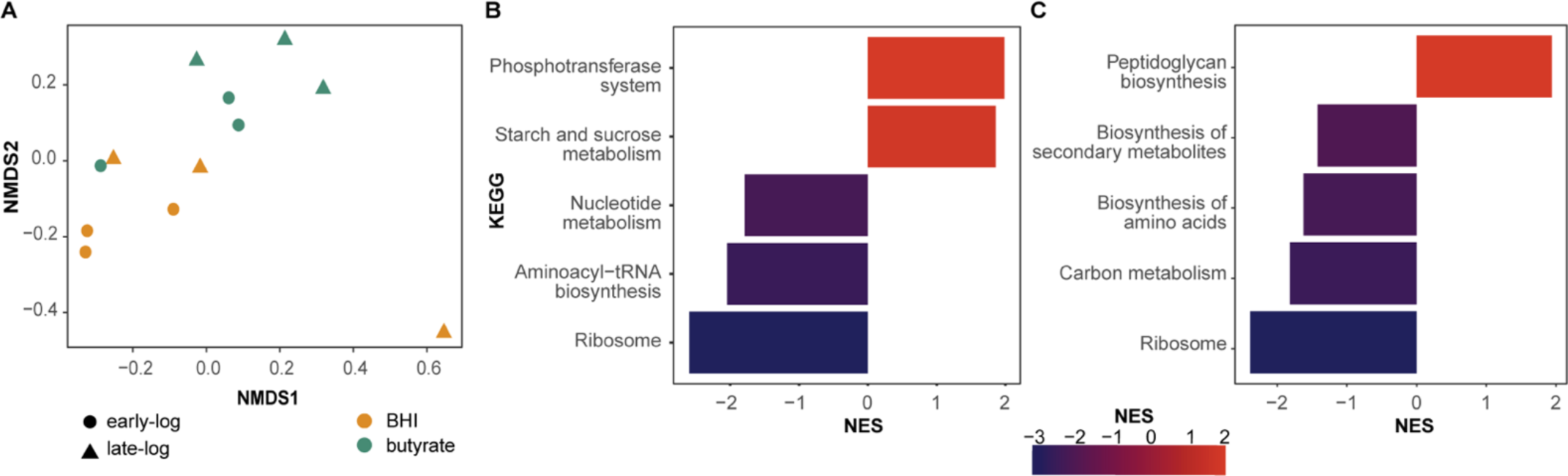
Butyrate modulates *C. difficile* gene expression. **A)** Non-metric multidimensional scaling (NMDS) of *C. difficile* strain 630 transcriptomic sequences using Bray-Curtis dissimilarity and normalized enrichment scores (NES) for KEGG assignments significantly up- and down-regulated genes at **B)** early-log (∼ 0.2 OD_600_), and **C)** late-log (∼ 0.5 OD_600_) for *C. difficile* strain 630 grown in BHI with or without 25 mM butyrate. NES was calculated using clusterprofiler (gseKEGG) in R with p-values adjusted post-hoc using False Discovery Rate.

At the level of individual genes, 38 and 8 genes were significantly over- or under-expressed during early-log growth in the presence of butyrate (Figure 6A and 6C, Wald’s test, log_2_ fold change > 1 and adjusted p < 10^-6^). Many of these included genes related to sporulation, such as stage II, III or even V sporulation proteins, as well as spore endopeptidases, which are required for spore germination and produced during sporulation (Figure 6C). The toxin gene TcdA was significantly over-expressed (Wald’s test, adjusted p <0.05) during early-log growth but not late-log, validating earlier production of toxin in the presence of butyrate (Supplementary figure S4 and S5). Several sigma factors (sigma-E, F, and G) that are involved in transcription regulation of sporulation typically expressed during later log growth (64–66) were also overexpressed during early-log growth in the presence of butyrate (Figure 6C). Several genes expressed significantly differentially during early-log growth were also related to metabolism. Genes related to glycine metabolism, such as the bi-functional glycine dehydrogenase/aminomethyl transferase protein (*gcvTPA*), and glycine decarboxylase (*gcvPB*) were upregulated in the presence of butyrate (Figure 6C). Other upregulated genes included phosphotransferase (PTS) genes related to lactose (PTS system, lactose/cellobiose family IIBC), fructose (PTS system, fructose/mannitol family IIB), mannose (PTS system, mannose specific IIBC), and mannitol (PTS system, fructose/mannitol family IIB), which collectively aid in non-glucose related carbohydrate metabolism (Supplementary figure S5) (67). Genes related to butyrate metabolism were also downregulated in the presence of butyrate, such as gamma-aminobutyrate dehydratase gamma-aminobutyrate-dehydratase (*abfD*) and 4-hydroxybutyrate dehydrogenase (*4hbD*) (Figure 6C), both involved in the succinate to butanoate fermentation pathway (68, 69).

**Figure 6:**
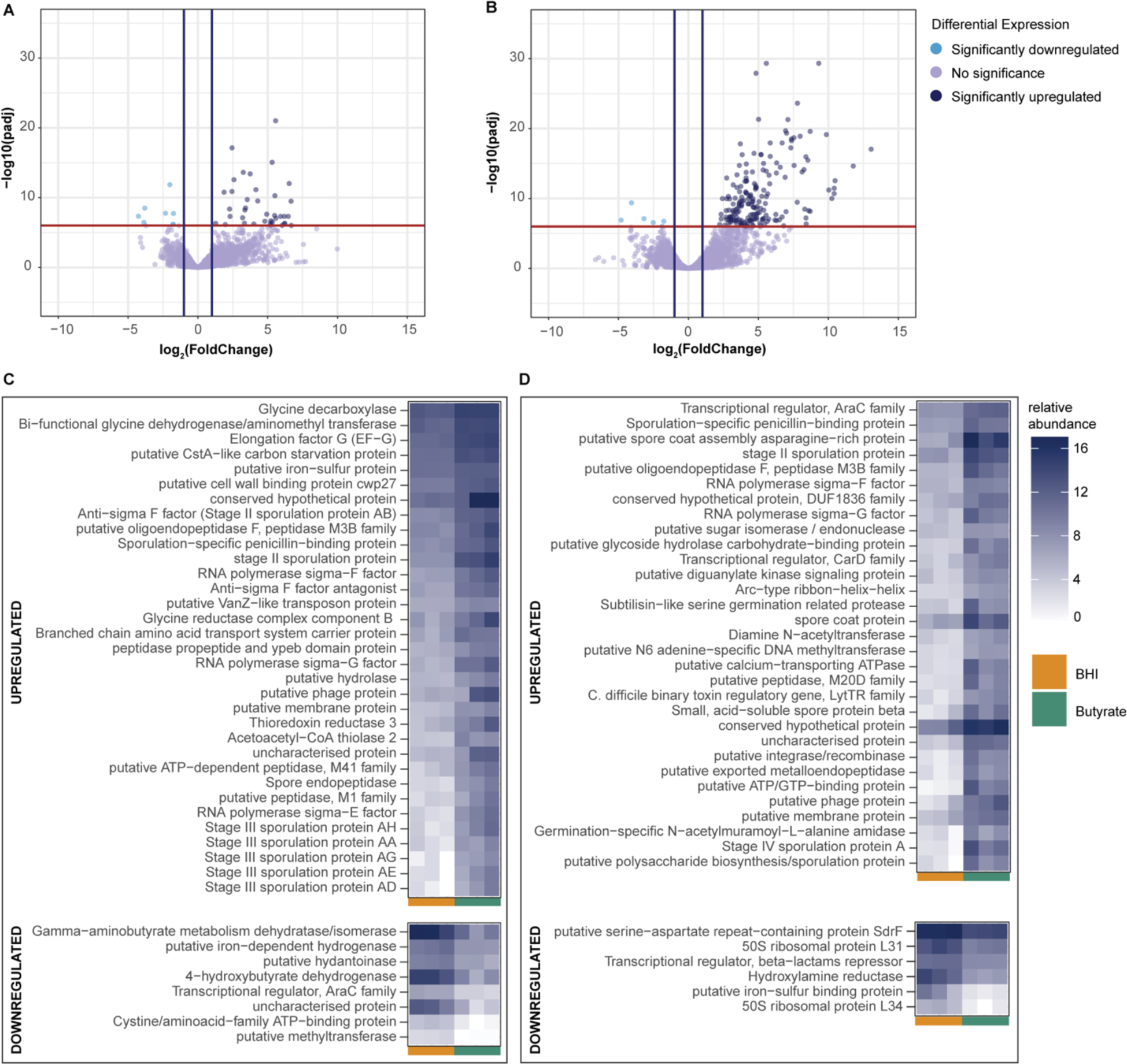
Butyrate upregulates genes related to spore formation and metabolism. Volcano plots of significantly upregulated and downregulated genes (Wald’s test, adjusted p < 10^-6^, log2foldchange > 1) at **A)** early-log and **B)** late-log growth of *C. difficile* strain 630 grown in BHI with or without 25 mM butyrate. Heatmap depicting normalized transcript counts of the top 50 significantly upregulated (top panels) and downregulated (bottom panels) genes in the presence of butyrate at **C)** early-log and **D)** late-log growth.

At late-log growth, 100 and 6 genes were over- and under-expressed, respectively (Figure 6B and 6D). Many sporulation related genes remained over-represented during later-log growth, including sporulation stage IV proteins, stage II sporulation proteins, subtilisin-like proteases, oligoendopeptidases and others, very likely due to the late-log also being the stationary phase where sporulation is observed *en masse*. Butyrate metabolism-associated genes remained downregulated in the presence of butyrate, gamma-aminobutyrate-dehydrogenase, and 4-hydroxy dehydrogenase. Most notably, genes associated with sporulation were upregulated in the presence of butyrate, many of which are involved in later steps of sporulation, such as spore coat and maturation proteins (spore coat proteins *sipL, cotB*; spore maturation proteins *spmA, spmB*; stage V sporulation proteins *spoIIIAB, spoIIIAC, spoIIIAF*). The binary toxin regulatory gene from LytR family of protiens, *cdtR*, was also upregulated. Other significantly downregulated genes included the ribosomal proteins L31 andL34, as well as a putative iron-sulphur binding protein, and genes related to antibiotic stress, including a transcriptional repressor for the beta-lactams, and hydroxylamine reductase, which is upregulated in response to metronidazole and fidaxomicin stress (70, 71).

We also investigated the involvement of known *C. difficile* global regulators in butyrate-induced growth changes, such as CcpA, Rex, PrdR, and CodY, which are known modulators of *C. difficile* virulence in response to its environment (11). None of these global regulators were significantly differentially expressed (Supplementary figure S4, S5). Additionally, we did identify over-representation of a putative carbon starvation protein, CstA in early-log (Figure 6C, S4, S5), and a histidine-kinase of *spo0A*, CD630_15790, was significantly upregulated in late log (Supplementary figure S4), which was recently identified to inhibit the activity of Spo0A (72).

## DISCUSSION

Butyrate has shown major promise in alleviating prominent intestinal diseases, such as graft-versus-host disease (73) or inflammatory bowel disease (74). In the context of CDI, higher butyrate levels are correlated with successful FMT in human studies (22) and inversely correlated with *C. difficile* burden in mice (75). While recent studies, including the data presented here, have demonstrated an inhibitory effect on *in vitro* growth of *C. difficile* (57, 75), the mechanism by which butyrate could inhibit *C. difficile* remains unknown. Our current results suggest a more complex role for butyrate in directly influencing *C. difficile*. Indeed, exogenous butyrate supplementation, while capable of attenuating disease via host effects, has not demonstrably reduced *C. difficile* burden in infected mice (25). Furthermore, recent studies in mice also suggest that the presence of butyrate-producing bacteria alone is not sufficient to inhibit *C. difficile* colonization (76).

Our results support previous observations that butyrate can inhibit growth of *C. difficile*. However, the ability of butyrate to inhibit *C. difficile* was not observed for all strains tested in our study, nor was it universally observed across different media. In contrast to a recent study observing butyrate-induced growth inhibition of various clinical strains (57), we observed limited inhibition by butyrate against two commonly used lab strains, *C. difficile* strain VPI10463 and R20291. For strain 630, growth inhibition by butyrate was also context-dependent; when grown *in vitro* with CDMM and a single carbohydrate source, *C. difficile* growth was significantly inhibited by butyrate only in the presence of raffinose and lactose. Other tested sugars, including mannitol, fructose, and mannose, demonstrated increased growth of *C. difficile* with butyrate compared to without. These observations make sense in an *in vivo* context, where a diverse milieu of metabolites and energy sources could possibly negate the inhibitory effects of butyrate. For instance, we also observed increased growth of *C. difficile in vitro* in the presence of acetate, another prominent SCFA in the gut, although these differences were not significant.

Our results also demonstrate the ability of butyrate to modulate *C. difficile* pathogenesis via spore and toxin production. To our knowledge, increased spore production by butyrate has not been previously demonstrated, although higher spore and toxin production has been predicted in response to increasing SCFAs, which were shown to decrease biofilm production (77). A previous study reported enhanced toxin production by *C. difficile* in the presence of butyrate, similar to our current study (78). This study also observed a correlation between toxin and butyrate production by *C. difficile* itself, whereby the addition of different amino acids downregulated the production of both. Furthermore, toxin production has been correlated with increased expression of butyrate metabolism in *C. difficile* in subsequent studies (79). This contrasts with our results, where we observed higher toxin production in the presence of butyrate, yet downregulation of 3-hydroxybutyryl-CoA dehydrogenase, an enzyme known to be involved in the production of butyrate and butanol in *Clostridium acetobutylicum* (80). Although these results might initially seem contradictory, we take these observations as further evidence for coordination of metabolism and toxin production by *C. difficile*, whereby the presence of butyrate might initially increase toxin production but later downregulate its production as the population of metabolizing cells are pushed into sporulation.

Perhaps more importantly, both our phenotypic and RNAseq data demonstrated a significant increase in *C. difficile* spore production in the presence of butyrate, which is connected to toxin production and metabolism (46). Indeed, a recent study demonstrated higher spore counts and increased disease severity in mice mono-colonized with a butyrate-producing bacterium, *Clostridium sardiniense*, prior to *C. difficile* infection (76). This is in contrast to impeded growth and attenuated disease in mice mono-colonized with *Paraclostridium bifermentans*, which can compete for amino acids via Stickland fermentation. Interestingly, our results mimic *in vivo C. difficile* RNAseq profiles of mice infected with pathogenic strains compared to strains deficient in toxin (30, 81), whereby PTS transport of alternative carbohydrate metabolic pathways, such as mannose, lactose or fructose, are preferred instead of glucose-focused pathways or other alternate carbon sources. Our RNAseq data in rich media also matches what we observed phenotypically in the presence of carbohydrate supplementation of CDMM, in which butyrate only had a positive growth impact in the presence of certain carbohydrates, such as mannose, lactose or fructose, reflected by the increased expression of PTS transporters of these carbohydrates in our RNAseq data. Though our data also shows an increase in expression of PTS transporters of mannitol, cellobiose and xylose, we did not test these carbohydrates due to an expectation that these will also show similar results to mannose, lactose and fructose *in vitro*.

In terms of how butyrate may impact regulation of *C. difficile* virulence (72), we might expect decreased *codY*, *ccpA* and *rex* expression in the presence of butyrate, given that we observed increased toxin in the presence of butyrate. CodY and CcpA typically decrease toxin and butyrate production (82, 83), whereas Rex is an important global regulator that responds to NAD^+^/NADH ratios in the cell, particularly when glucose or other rapidly metabolized sugars are not around (11, 84, 85). While we observed increased *ccpA* expression during early- and late-log growth from our qPCR (∼0.8 mean logFoldChange) but significantly decreased in our RNAseq (-1.23 logFoldChange) only in early-log, our results from both RNAseq and qRT-PCR for the canonical virulence regulators (*codY, rex, prdR*) were not significant in the presence of butyrate. The involvement of these genes cannot be resolved from our data alone, and it is possible that regulatory responses to butyrate may be independent of these regulators, even though our data suggests a metabolic connection. Interestingly, we observed significant upregulation of a putative carbon-starvation gene, *cstA*, during early-log growth. The canonical gene *cstA* has been demonstrated to be involved in peptide utilization, agglutination and motility in another gut pathogen, *Campylobacter jejuni* (86), which was not upregulated in our data. Given the effect of butyrate on toxin and spore production, as well as the types of genes that were upregulated by it, it is possible that butyrate induces a collective stress response via alternate regulatory mechanisms leading to premature induction of sporulation in vegetative cells.

Independent of the potential regulatory mechanisms, the effect of butyrate on *C. difficile* has important clinical implications (Figure 7). While butyrate and *C. difficile* levels are consistently negatively regulated following successful FMT and have been demonstrated to attenuate inflammation via the host, our results suggest a complicated outcome for butyrate-focused treatments. For instance, treating patients with CDI with butyrate alone (either with butyrate-producing bacteria or exogenous application) may have a detrimental effect on the patient, as has been observed *in vivo* in mice (76).Yet, combining this approach with additional microbiota members that can compete with *C. difficile* for nutrients may appropriately supplement the anti-inflammatory, and potentially inhibitory, effect by butyrate in the gut environment. Identification of the regulatory elements that dictate the effects of butyrate may expedite these findings.

**Figure 7:**
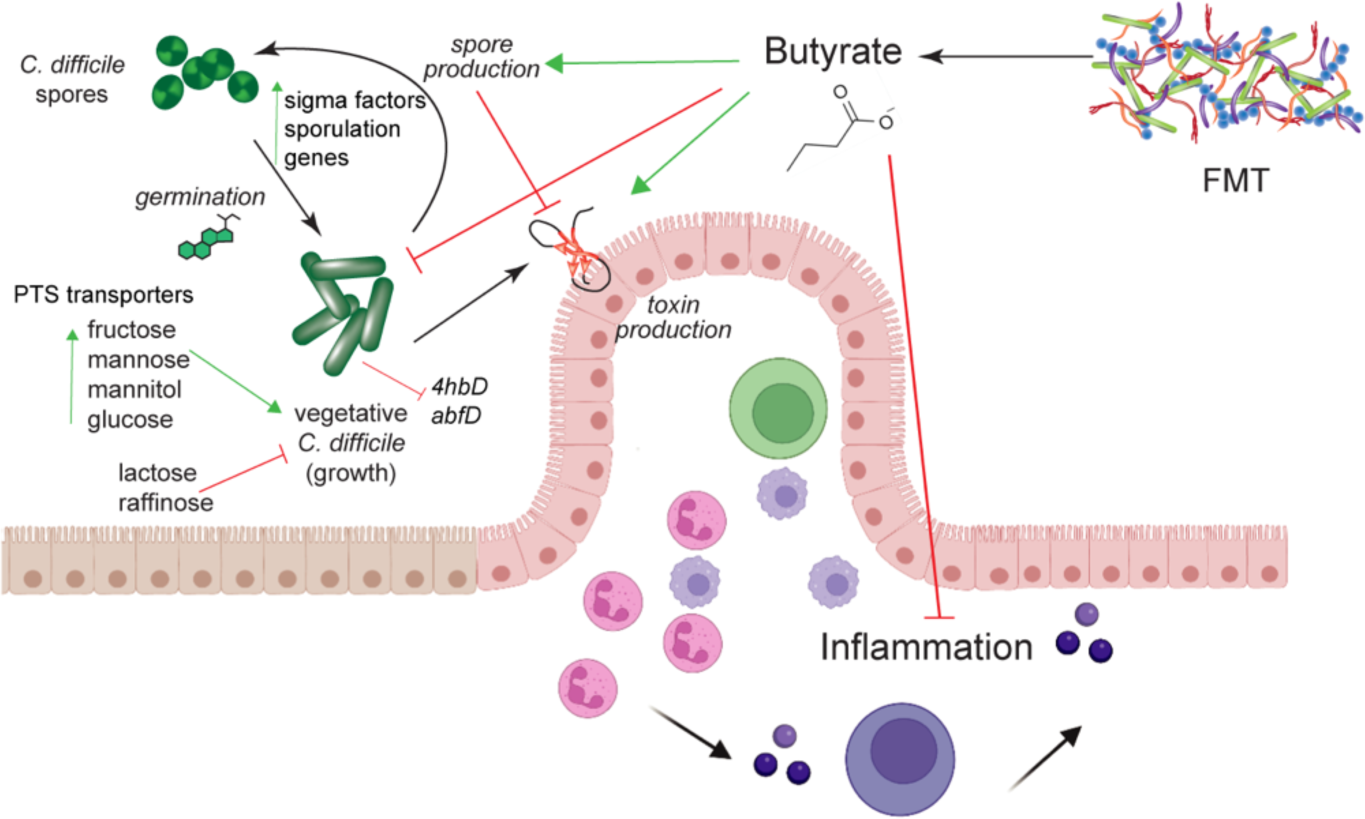
Model for potential mechanism of butyrate effectiveness against *C. difficile* strain 630. Increasing butyrate may alleviate host inflammation during recovery of the microbiota (such as via fecal microbiota transplantation, or FMT) but also signals *C. difficile* to change metabolic strategies to increase survival. This may involve increased expression of PTS transporters for mannose, fructose, and mannitol, inducing alternate metabolic pathways for carbohydrate utilization in metabolically active cells. While growth of vegetative cells may be inhibited, sporulation and toxin genes are upregulated to optimize colonization. Figure illustrated in part with Biorender.

## ACKNOWLEDGEMENTS

We would like to acknowledge Clemson University for generous allotment of compute time on Palmetto cluster. This publication was made possible, in part, with support from the Clemson University Genomics and Bioinformatics Facility, which receives support from an Institutional Development Award (IDeA) from the National Institute of General Medical Sciences of the National Institutes of Health under grant number P20GM109094. AMS was supported by grant number K01-DK111794 from the National Institute of Diabetes and Digestive and Kidney Diseases. We would like to thank Clemson University Department of Biological Sciences Microbiology prep labs for the use of their phase contrast microscope. AMS has received consultation fees from Finch Therapeutics and Rebiotix / Ferring Pharmaceuticals.

M.B. – Conceptualization, Formal analysis, Data Curation, Investigation, Methodology, Validation, Writing—original draft; J.C. – Data Curation, Formal analysis, Investigation, Methodology, Validation, Writing – original draft; D.B. - Formal analysis, Methodology, Software, Writing—original draft, and Writing—review and editing; S.N. – Methodology, Investigation; A.M. – Methodology, Investigation; A.M.S. – Supervision, Project Administration, Funding Acquisition, Conceptualization, Writing – original draft, Writing – review and editing.

## Supplemental Outline

**Supplementary Table S1: RNA-seq preliminary statistics.** Preliminary raw read numbers and statistics generated in the process of analysis.

**Supplementary Figure S1. Butyrate does not significantly impact the growth of *C. difficile* strains R20291 or VPI10643. A)** Growth curves measuring OD_600_ of *C. difficile* R20291 over 24 hours in presence of 5 and 25 mM acetate, propionate, and butyrate in BHI. **B)** Growth curves measuring OD_600_ of *C. difficile* VPI10463 over 24 hours in presence of 5 and 25 mM acetate, propionate, and butyrate in BHI.

**Supplementary Figure S2. Butyrate-induced growth inhibition is not impacted by pH.** Growth of *C. difficile* strain 630 over 24 hours at pH 6.2 (panel 1), 7.2 (panel 2), and 8 (panel 3) in BHI supplemented with 5 or 25 mM acetate, propionate, and butyrate. P-value through Dunnett’s test, *p-value < 0.05; **p-value < 0.01; ***p-value < 0.001.

**Supplementary Figure S3. Short chain fatty acids increase toxin production in *C. difficile* strains R20291 and VPI10463.** Toxin activity of *C. difficile* strain **A**) R20291 and **B)** VPI10463 over 24 hours in BHI supplemented with 5 or 25 mM acetate, propionate, and butyrate. **C)** Toxin activity *C.difficile* 630 at 6, 12, 18 and 24 hours in BHI supplemented with 5 or 25 mM acetate, propionate, and butyrate . P-value through Dunnett’s test, *p-value < 0.05; **p-value < 0.01; ***p-value < 0.001.

**Supplementary Figure S4. Butyrate increases the expression of carbohydrate metabolism genes.** Heatmap of selected significant genes representing the log2FoldChange of growth in butyrate versus without butyrate in early- and late-log calculated using DESeq in R as described for RNASeq analysis in Materials and Methods. Significance (p <0.05) was calculated using Wald’s test; all genes listed are significantly over- or under-expressed over growth in BHI alone except *ccpA* (not significant).

**Supplementary figure S5. Butyrate modulates select *C. difficile* carbohydrate metabolism and virulence regulators.** Relative abundance of normalized transcript counts of select significant genes in early- and late-log. Normalized transcripts were obtained using nnTransform in DESeq. The significance was calculated by one-way ANOVA between transcripts from BHI growth with or without butyrate *p-value < 0.05; **p-value < 0.01; ***p-value < 0.001

